# Processing of Genomic RNAs by Dicer in Bat Cells Limits SARS-CoV-2 Replication

**DOI:** 10.1101/2025.01.10.632425

**Authors:** Iyanuoluwani J. Owolabi, Shazeed-UI Karim, Sweta Khanal, Sergio Valdivia, Christopher Frenzel, Fengwei Bai, Alex S. Flynt

## Abstract

Bats are reservoirs for numerous viruses that cause serious diseases in other animals and humans. Several mechanisms are proposed to contribute to the tolerance of bats to these pathogens. This study investigates the response of bat cells to double-stranded RNA generated by SARS-CoV-2 replication. Here, we found the involvement of Dicer in the processing of viral genomic RNAs during SARS-CoV-2 infection. Examining RNA sequencing of infected cells, small-interfering RNA (siRNA)-like fragments were found derived from viral RNAs. Depletion of Dicer showed a reduction in these RNAs and an increase in viral loads suggesting unlike other mammals, bats may use Dicer to limit viral replication. This prompted the exploration of key dsRNA sensors in bat cells. Our analysis showed significant upregulation of OAS1 and MX1 in response to dsRNA, while PKR levels remained low, suggesting alternative dsRNA-response mechanisms are present that eschew the common PKR-based system. These results further show how bats employ distinct strategies for antiviral defense that may contribute to tolerating viral infections. They suggest the involvement of Dicer in antiviral mechanisms in bats, a function not observed in other mammals. This highlights a mechanism for bat originating viruses to evolve features that in other animals could cause extreme antiviral responses such as is seen with SARS-CoV-2.

**GRAPHICAL SUMMARY:** 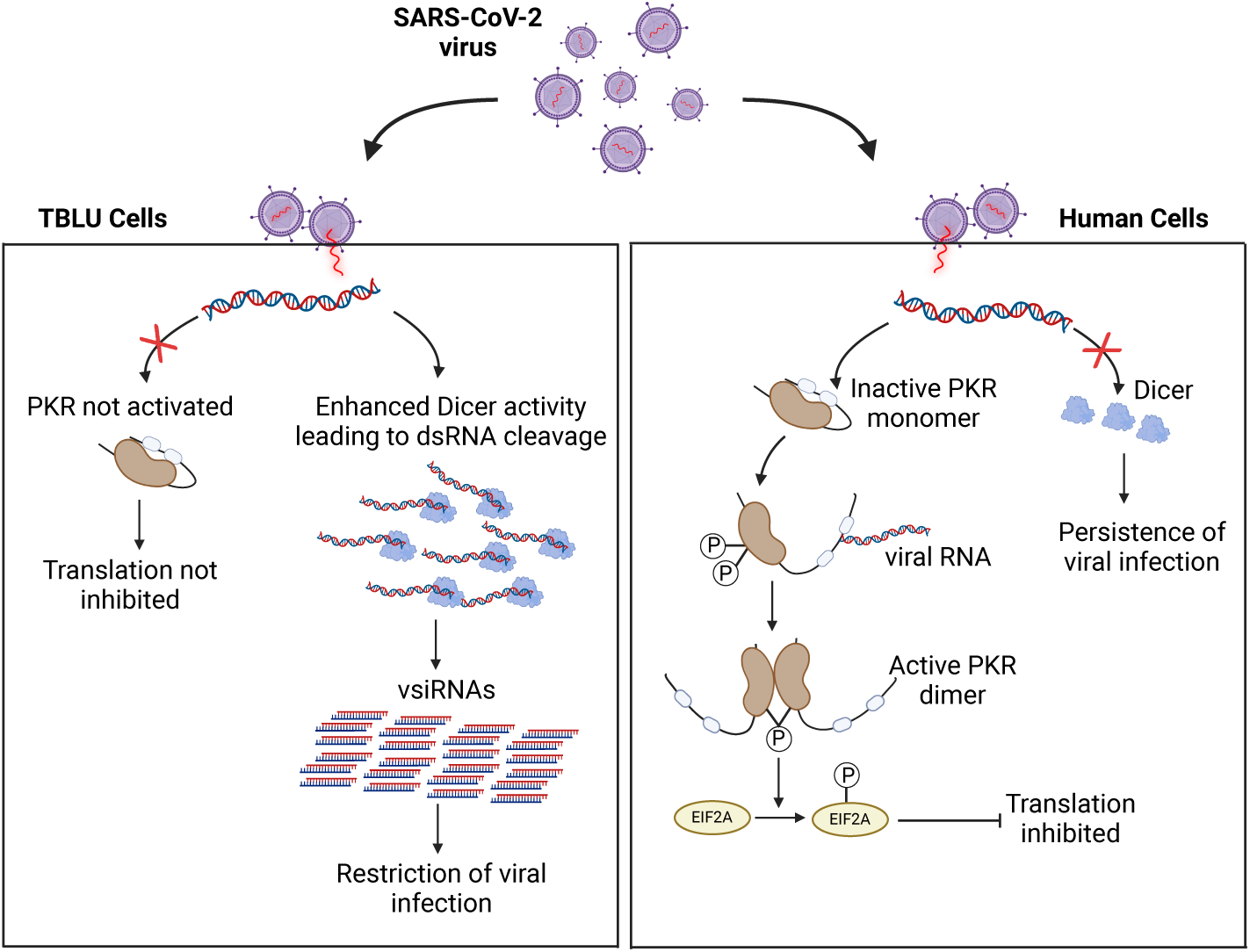

## BACKGROUND

Viruses are common in bats, including 4,800 coronaviruses which make up 30% of all known bat viruses (1). Bats are as the reservoir for lineages of betacoronaviruses, from which several human pathogens have evolved, such as SARS-CoV, MERS-CoV, and SARS-CoV-2 (2,3). This study focuses on SARS-CoV-2, which emerged in late 2019 in Wuhan, China, and shares 79% and 50% sequence similarity with SARS-CoV and MERS-CoV, respectively (4). The SARS virus primarily enters host cells by binding to the angiotensin-converting enzyme 2 (ACE2) receptor (2,5), which is expressed in various tissues, including the nasopharynx, lungs, intestines, and heart (6–8). SARS-CoV-2 has a ∼30 kb genome encoding structural proteins like Spike (S), Envelope (E), Matrix (M), and Nucleocapsid (N), which facilitate viral entry, assembly and RNA packaging, along with non-structural proteins such as nsp3 and nsp12 that are crucial for genome replication (5,9). Bats are also the origin of the two oldest human coronaviruses, HCoV-229E and HCoV-NL63, which cause the common cold (10,11)^-^. Beyond coronaviruses, bats harbor other high-priority zoonotic viruses such as Rabies virus, Nipah virus, Hendra virus, Ebola virus, and Marburg virus, all of which have raised substantial global public health concerns (12–15).

There are many hypotheses about why bats are the reservoir for many viruses without causing diseases in themselves. For one, during flight, bat body temperature increases and resembles the febrile response in other mammals. This may exert a selective pressure on viral pathogens to persist at elevated body temperatures (16). The proportional increase in body temperature during viral infection observed in bats is trivial compared with the rise during flight (16). In addition to unique flight physiology, there are many changes to the bat immune system. The overall percentage of immune-related genes in the bat genome is significantly lower compared to the 7% observed in humans (17). Bat species such as *Pteropus alecto (P. alecto)*, *Artibeus jamaicensis*, and *Rousettus aegyptiacus* possess only 3.5%, 3.26%, and 2.75% immune-related genes, respectively (18–20). Known immune modifications in bats include the suppression of type I and type II interferon (IFN) production, regulation of inflammation through the modulation of NF-κB signaling, and inhibition of NLRP3 inflammasome activation during viral infections (21,22). An apparent consequence of the innate immune changes is that bats exhibit less intense and shorter-lived adaptive immune responses (23,24), suggesting immune responses have evolved to tolerate and coexist with viruses rather than mounting aggressive responses for viral elimination.

This reconfigured response to viral infection extends to bat pattern recognition receptors (PRRs) like Toll-like receptors (TLRs) that have evolved key amino acid differences in ligand-binding domains (25). Bats show positive selection in TLR3, TLR7, TLR8, and TLR9, with selection concentrated in the leucine-rich repeat domains involved in pathogen recognition (21,26,27). Additionally, bats lack the PYHIN PRR gene family, which diminishes their DNA-sensing capability (25,28). This fine-tuning of PRR signaling results in reduced immune responses, including lower interferon production and minimal inflammation (21,22,29).

Given that anti-viral responses in bats are dampened, this raises the possibility that dsRNA generated during viral replication may also elicit a distinct response. In most mammals, RNA-sensing receptors like PKR, OAS1, and MX1 recognize the biochemical characteristics of dsRNA (30). In bats, exposure to synthetic dsRNA leads to the rapid activation of OAS1 and MX1, followed shortly by attenuation of expression (21,31). In contrast, human cells sustain elevated levels of these RNA sensors for a significantly longer period, extending over 16 hours beyond the duration observed in bat RNA sensors in response to dsRNA (31). Another critical dsRNA sensor, PKR, does not show significant upregulation in bats (32). One possible explanation for the reduced PKR response is to avoid inducing translational arrest through PKR-mediated phosphorylation of eIF2α, which activates stress-related genes such as ATF3 and ATF4 (33). If not carefully regulated, this activation could lead to cellular damage. The diminished PKR response in bats may suggest that cytoplasmic dsRNA is not prohibited and raises the question as to how these molecules are otherwise cleared from cells.

Dicer, a dsRNA processing enzyme, has antiviral activity in many eukaryotes, such as insects, through destruction of viral replication intermediates (34–36). Invertebrate Dicer can process viral RNA into virus-derived small-interfering RNAs (vsiRNA), which can be incorporated into effector complexes for RNA silencing (37). Additionally, Dicer’s processing of viral RNA alone can also limit replication (36). The effectiveness of Dicer in eliminating viral RNAs likely explains why many viruses have evolved anti-Dicer strategies. These strategies often involve the production of viral suppressors of RNAi (VSRs), which inhibit Dicer activity through mechanisms such as sequestering dsRNA or vsiRNAs to block Dicer processing (38). Beyond interfering with antiviral RNAi, VSRs may also disrupt miRNA- and endo-siRNA-mediated gene silencing, further compromising host cellular regulation (39–42) In contrast, vertebrates have apparently abandoned Dicer and small RNA-mediated viral immunity in favor of cell-mediated and innate immune responses in response to viral replication. Considering the changes in bat dsRNA sensing, this raises the possibility of a reactivated role for Dicer in processing viral RNAs.

Here, we explore the role of Dicer in the antiviral response mechanism in bat *Tadarida brasiliensis* (*T. brasiliensis*) lung cells (TBLU) to dsRNA generated during SARS-CoV-2 replication. Our findings reveal that 22 nt siRNA-like RNAs are generated by Dicer from viral genomic RNAs. Depletion of Dicer resulted in a reduction of these siRNA-like RNAs, suggesting that, unlike other mammals, bats may utilize Dicer to limit viral replication. Furthermore, our analysis demonstrated that PKR levels remained low during SARS-CoV-2 infection, suggesting the existence of alternative dsRNA-sensing mechanisms applies to this virus. Collectively, these results indicate that bats may use Dicer in managing SARS-CoV-2 dsRNA accumulation and contributing to the tolerance of infection.

## RESULTS

### Establishing a TBLU cell line permissive to SARS-CoV-2 infection

TBLU and Vero E6 cells were infected with SARS-CoV-2 at a multiplicity of infection (MOI) of 0.5 and 1.0. At 48 hours post-infection, SARS-CoV-2 nucleocapsid RNA was detectable in infected Vero E6 cells but not in TBLU cells (Figure 1A) (Supp Figure 1), consistent with reports that ACE2 is lost in the TBLU(43). To create TBLU cells permissive to SARS-CoV-2 infection, we generated a line stably expressing human ACE2 (TBLU-ACE2). Immunostaining and qPCR for the SARS-CoV-2 nucleocapsid protein confirmed infection in TBLU-ACE2 cells, with nucleocapsid protein and transcripts readily detected in both TBLU-ACE2 and VeroE6 cells (Figure 1A). Notably, Vero E6 cells exhibited a higher viral load than TBLU-ACE2 cells, suggesting either increased susceptibility to infection or more robust viral replication (Figure 1B). To further assess viral replication and confirm the replication of SARS-CoV-2 genome was taking place in the infected bat cells, we measured the presence of the sub-genomic RNAs (sgRNAs) encoding the key structural proteins of the virus, including spike (S), matrix (M), nucleoprotein (N), and envelope (E). All sgRNAs were amplified, verifying progression through the viral life cycle (Figure 1C). The detection of these sgRNAs suggests that SARS-CoV-2 not only enters and infects TBLU-ACE2 cells but also undergoes productive replication and therefore, generates dsRNA.

**Figure 1.**
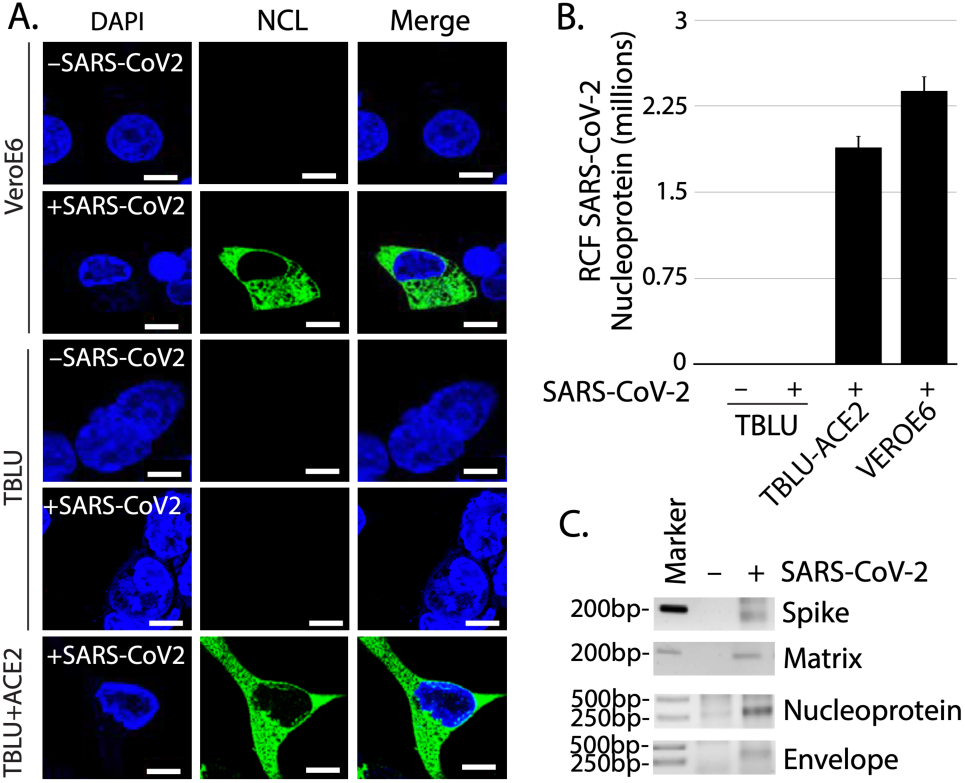
Generation of a permissive TBLU cell line to SARS-CoV-2 infection. **A)** Immunostaining analysis of SARS-CoV-2 nucleocapsid protein expression in Vero E6, TBLU, and TBLU-ACE2 cells 48 hours post-infection (0.5 MOI). **B)** Quantification of SARS-CoV-2 nucleocapsid viral expression in TBLU, TBLU-ACE2, and Vero E6 cells. Different letters denote significance determined by Tukey ANOVA (p ≤ 0.05). (C) Detection of SARS-CoV-2 sgRNAs encoding spike (S), matrix (M), nucleoprotein (N), and envelope (E) proteins in TBLU-ACE2 cells.

### Dicer knockdown leads to greater viral replication and decreases vsiRNA accumulation

Using these cells, we then used an RNAi approach to investigate whether loss of Dicer impacted SARS-CoV-2 infection in the TBLU-ACE2 cells. We achieved 50% depletion of Dicer in Dicer knockdown (DKD) TBLU-ACE2 cells (Figure 2A). DKD and control TBLU-ACE2 cells were infected with SARS-CoV-2 at the same MOI. Viral replication was significantly higher in DKD cells compared to control cells (Figure 2B). To compare small RNA profiles between the two conditions, we concatenated all small RNA libraries from infected control TBLU-ACE2 cells into a control dataset and the DKD small RNA data into a DKD dataset. Reads in the control library had a mapping rate of 0.12% to the SARS-CoV-2 genome, while the DKD library had a lower mapping rate of 0.07%. The lower mapping rate in DKD could result from less fragmentation of viral RNA and slower turnover of genomic RNAs in the absence of Dicer. Examining the mapping of sense and antisense vsiRNAs in the two conditions revealed no difference in the abundance of 17–19 nt small RNAs derived from SARS-CoV-2 between the control and DKD libraries for both mapping orientations (Figure 2C-D). The 17–19 nt small RNAs are likely degradation products as their size falls below the typical Dicer product range (20-24nt). There is no substantial difference in the abundance of 20–24 nt sense vsiRNAs between the two conditions (Figure 2E-F). In contrast, we observe a greater abundance of antisense 20–24 nt vsiRNAs in the control library compared to the DKD library (Figure 2E-F). SARS-CoV-2 antisense RNAs are replication templates and thus are predominantly dsRNA, unlike sense RNAs. The loss of the antisense small RNAs in the Dicer size range (20-24nt) following Dicer knockdown suggests the fragments could be *bona fide* vsiRNAs.

**Figure 2.**
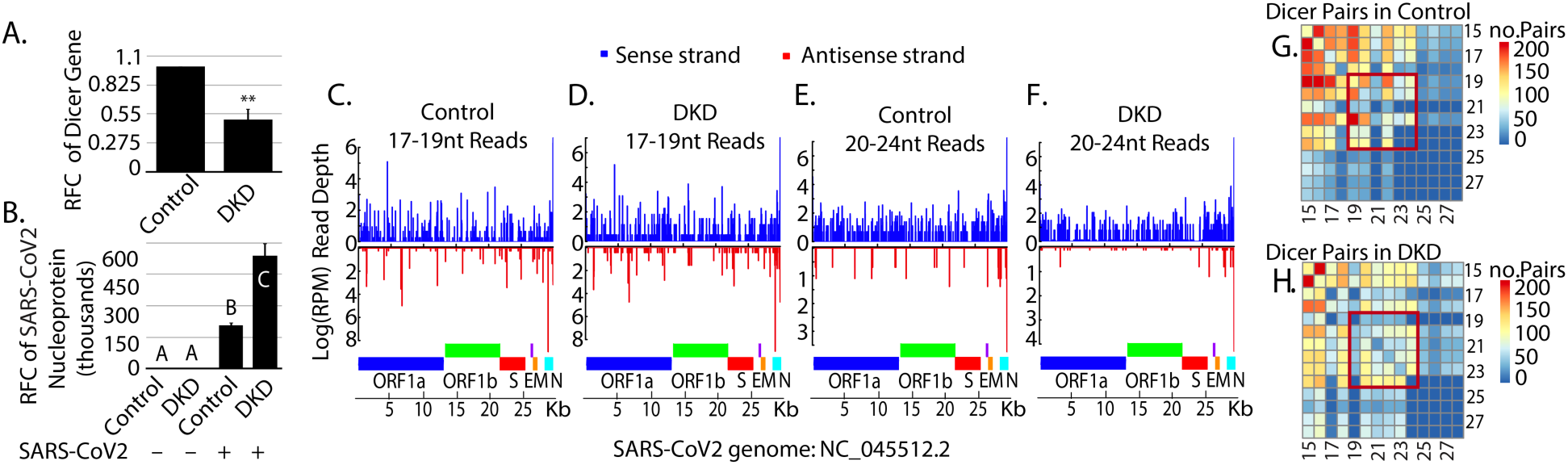
Antiviral Role of Dicer in SARS-CoV-2-Infected TBLU-ACE2 Cells. **A)** Quantification of Dicer expression in control and DKD cells by qRT-PCR. Control showed a significantly higher value compared to DKD (P-value = 0.002, t-test), **B)** Bar graph depicting SARS-CoV-2 nucleocapsid protein expression in control and DKD cells. In both bar graphs, the different letters denote significance determined by Tukey ANOVA (p ≤ 0.05) **C-F)** Sushi plots illustrating the mapping of sense (blue) and antisense (red) small RNA sequencing reads to the SARS-CoV-2 genome in control and DKD cells. Major genomic features of the genome are underneath as colored bars. **C-D)** Mapping of 17-19 nt reads is measured, and in **(E-F)** 20-24 nt reads. **G-H)** Heatmap representing overlapping pairs of reads based on mapping to the SARS-CoV-2 genome. The size of the reads that form pairs are shown at the bottom and right of each heatmap. The red box highlights the sizes of RNAs expected for Dicer products.

To further determine if the potential vsiRNAs are Dicer products, we quantified overlapping read pairs that exhibit 2-nucleotide 3’ overhangs, which is characteristic of Dicer processing. The numbers of Dicer pairs of different lengths were then visualized in a matrix showing 15–31 nt (rows) RNAs paired with 15–31 nt small RNAs (columns) (Figure 2G-H). In the control dataset, hundreds of 20-24 nt, nearly symmetrical pairs were observed (Figure 2G). In the DKD condition pairs that exhibit the characteristics of siRNAs (20-24nt long and symmetrical) were significantly decreased (Figure 2H). Small RNA pairs outside this size range are asymmetrical pairs that are unlikely to be Dicer products. Our findings are consistent with previous studies showing that in contrast to mammalian cell lines like Vero E6, 293T (cells derived from human embryonic kidney cells SV40 T-antigen), and BHK (fibroblast cells derived from the kidney of hamsters), which failed to generate vsiRNAs after Sindbis virus (SINV) infection (44–46), the presence of vsiRNAs was observed in SINV-infected bat-derived PaKi cells (kidney cells derived from *P. alecto*) (47). These differences underscore the bat specific activity of Dicer in processing viral dsRNA into siRNA, and our study includes SARS-CoV-2.

### SARS-CoV-2 Infection Remodels Host Small RNA Profiles

The interaction of Dicer with viral RNAs led us to investigate how SARS-CoV-2 infection may influence the processing of endogenous small RNAs. To do this, we first identified genomic loci that give rise to small RNAs based on depth (>100 reads) from small RNA sequencing alignment as well as a minimum length (>60 bp). Calling loci with these parameters should capture microRNAs (miRNAs) and other classes that are derived from longer precursors. This led to annotation of 3530 loci in the *T. brasiliensis* genome (ACC#GCA_004025005.1) (Supplementary file 1). Change in small RNA expression was then assessed by comparing libraries from infected and uninfected TBLU-ACE2 cells, which showed a dramatic perturbation in endogenous small RNA expression, where ∼68% of loci were differentially expressed based on p-value ≤ 0.05 (Figure 3A) (Supplementary file 2). Altered expression was even more evident for loci with a >80% bias for reads in the 20-24nt range–the sizes expected for Dicer processed RNAs. Approximately 84% of putative Dicer processed loci were differentially expressed after SARS-CoV-2 infection.

**Figure 3.**
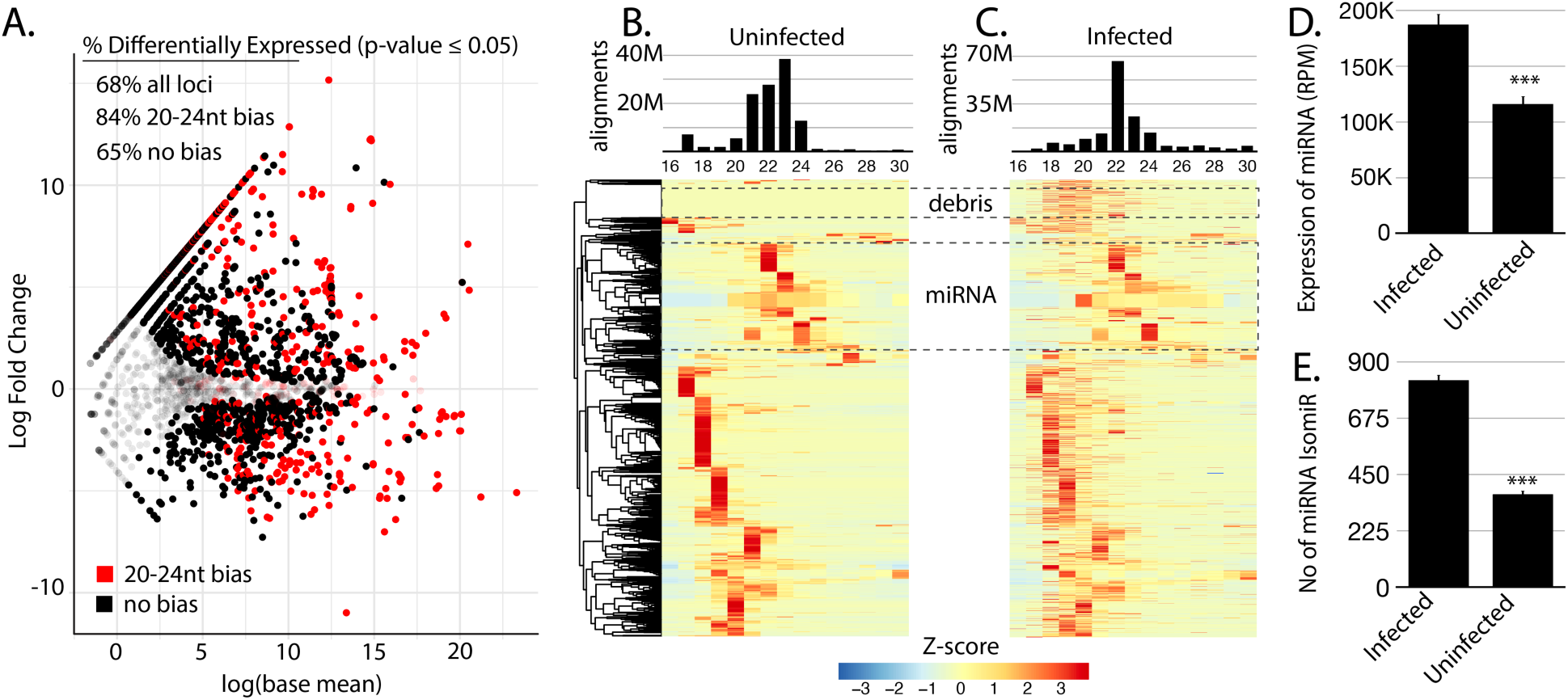
Altered host small RNA expression after SARS-CoV-2-Infection in TBLU-ACE2 Cells. **A)** MA plot showing differential expression of small RNAs annotated in this study comparing infected to uninfected TBLU-ACE2 cells. Transparency was applied to small RNA species that did not show differential expression based on P-value ≤ 0.05. Small RNA-generating loci with a greater 80% bias for 20-24nt RNAs (potential Dicer products) are colored in red. **B-C)** Heatmap showing clustering of small RNAs based on the distribution of read size mapping to small RNA expression loci. Each row represents a different small RNA locus. Clustering was performed using RNA read sizes from uninfected cells **(B),** while the order of the infected heatmap **(C)** is based on the order in part B. Dashed lines show regions of interest. Debris highlights RNAs only seen after infection. The miRNA box designates the location of loci where mapping of Dicer product-sized RNAs occurs. **D)** Global miRNA expression in SARS-CoV-2-infected cells and uninfected cells. **E)** Number of total miRNA isomiRs expressed in infected and uninfected cells. Statistical significance for panels D and E was determined using a t-test, with p-values of 0.0006 and 9.72 × 10⁻¹², respectively.

Next, we analyzed the distribution of read lengths that map to each locus to further characterize changes in small RNAs after infection (Figure 3B-C). In both infected and uninfected datasets, Dicer products (miRNAs and siRNAs) appear to be the most abundant species based on the dominant 20-24 nt peak in the overall smallRNA population. However, the size distributions varied with infected library showed a peak at 22 nt, while uninfected library exhibited a peak at 23 nt (Figure 3B-C). Futher changes in expression were evident when examining size distribution per locus. Hierarchical clustering of loci based on the sizes of reads in the uninfected condition (Figure 3B) showed there were shifts in sizes of reads mapping to the same loci after infection (Figure 3C). This was most apparent in a group of loci that seem to be degradation fragments we are denoting as debris due to read size and heterogeneity that only show expression in the infected condition. This suggests greater fragmentation of host RNAs may occur during viral infection. The other group is the loci that have predominant 20-24nt mapping, which contains miRNAs. Here small, yet clear shifts in read size bias were seen in this group of RNAs in the infected group. Together these results show that not only are small RNAs differentially expressed after SARS-CoV-2 infection, but that there is a perturbation of biogenesis.

To focus specifically on Dicer products, we identified miRNA loci in *T. brasiliensis.* First, using sequencing data we annotated miRNAs in the *T. brasilensis* genome, finding 105 highly confident loci (Supplementary file 3). Comparing the expressions in the infected and uninfected revealed a nearly two-fold increase in the abundance of miRNA mapping reads (Figure 3D, Supplementary file 4). To also investigate the impact on the processing of miRNA, we assessed the diversity of reads mapping to miRNAs (Figure 3E). Unique sequences were identified to quantify the collection of miRNA isoforms (isomiRs) (Supplementary file 5). 827 isomiRs were found in the infected library compared to 370 in the uninfected library, indicating that not only was there an overall increase in miRNA expression during SARS-CoV-2 infection, but also less precision in the biogenesis of miRNAs (Figure 3D-E). These observations are consistent with a significant impact on Dicer activity by SARS-CoV-2 infection.

### Alternative Antiviral Response to SARS-CoV-2 infection in Bat Cells

Motivated by the apparent altered small RNA biogenesis observed in SARS-CoV-2-infected cells, we analyzed mRNA expression using public annotations (GCA_030848825 xenoRefGene) to assess changes in mRNA expression after infection of TBLU-ACE2 (Supplementary file 6). Of 23,562 mRNAs, 7,902 were differentially expressed based on p-value ≤ 0.05 (Figure 4A). Differentially expressed genes included Dicer which was upregulated by 3.38 log_2_ fold change. TLR3 was also upregulated by an even greater degree though with less statistical confidence, which suggests that viral dsRNA is present to trigger this change that occurs in responding to dsRNA(48). In contrast PKR, OAS3 and MX1 did not show any significant changes in response to infection indicating the virus did not generate sufficient dsRNA to trigger OAS3 or MX1, which are known to become upregulated in bats when exposed to dsRNA (32). This shows that while SARS-CoV-2 infection leads to the upregulated of Dicer and TLR3 expression, the response to dsRNA seen in other mammals does not occur in the bat cells in response to this virus.

**Figure 4.**
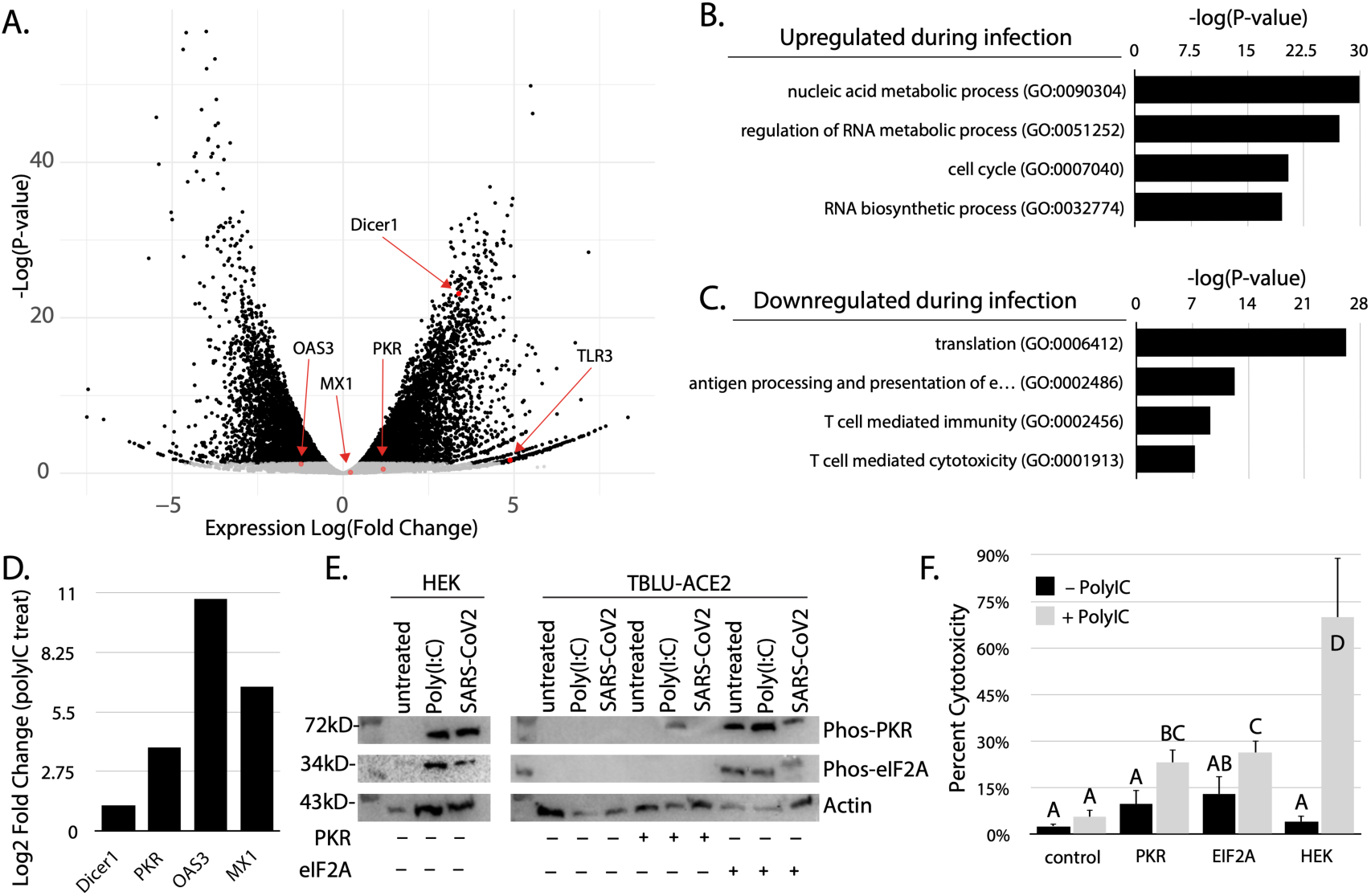
Antiviral responses in TBLU-ACE2 cells to viral infection and Poly(I:C) treatment. A) Differential Expression of mRNAs in TBLU-ACE2 cells after SARS-2-CoV-2 infection. Genes of interest are colored by red dots and designated with arrows and labels. Grey dots in the blot are not significantly different based on P-value ≤ 0.05. **B)** GO analysis of genes graphed in part A that were assigned a P-value ≤ 0.05 and upregulated greater than 2 log2 fold change. **C)** GO analysis of genes graphed in part A that were assigned a P-value ≤ 0.05 and downregulated less than -2 log2 fold change. **D)** Expression changes expressed as log2 fold change of select genes based on RNA sequencing after Poly(I:C) treatment in TBLU cells compared to untreated controls. **E)** Western blot analysis of phospho-PKR and phospho-eIF2α in Poly(I:C)-treated, or SARS-CoV-2 infected TBLU and HEK293 cells. TBLU cells were also transfected with plasmids encoding either PKR or eIF2α as shown by the plus and minus designations below the blot bands. **F)** LDH cytotoxicity assay measuring cell viability in Poly(I:C)-treated TBLU, HEK293, and PKR or eIF2α transfected bat cells. Statistical significance determined by Tukey ANOVA (P-value ≤ 0.05).

Further investigation of the differentially expressed genes using GO enrichment analysis found some unexpected biological processes occur in response to SARS-CoV-2 infection (Figure 4A-B). Upregulated genes, which we define as P ≤ 0.05 and > 2 log_2_ fold change, had multiple GO categories relating to RNA processing that were significantly enriched (Figure 4B) (Supplementary file 7). This indicates that in addition to Dicer expression changes, a variety of other genes also process RNA are expressed in response to SARS-CoV-2. Dicer is not upregulated in Humans in response to SARS-CoV-2 infection, suggesting that these changes in RNA metabolism may represent a novel antiviral mechanism specific to bats (49). Even more unexpected were the identity of downregulated genes– P ≤ 0.05 and < -2 log_2_ fold change (Figure 4C) (Supplementary file 8). We found that many anti-viral genes related to T-cells were decreased, including antigen presentation. The process of translation itself was also implicated, which invokes the primary outcome of PKR activation. It is unclear if these changes are driven by the virus or the host, but they show that bat cells have altered RNA-related processes but not activation of immune genes upon SARS-CoV-2 infection.

To verify that dsRNA sensing was intact in the TBLU-ACE2 cells, we exposed them to Poly(I:C) and measured the expression by RNA sequencing (Figure 4D). In a near reversal of the SARS-CoV-2 infection condition, Dicer was not upregulated, while PKR, OAS3, and MX1 were considerably upregulated. PKR had a modest expression increase from 7 FPKM to 105 FPKM when compared to OAS3, which increased from 0.89 FPKM to 1539 FPKM, and MX1, which went from 34 FPKM to 3443 FPKM. This is consistent with other studies that show OAS1 and MX1 expression increases in response to dsRNA in bats (32). Our observation that PKR is upregulated in TBLU cells could be a species-specific behavior. Nevertheless, the TBLU-ACE2 cells retained the expected cellular responses to dsRNA, which is not triggered by SARS-CoV-2 infection.

Next, we examined the post-translation modifications associated with PKR activation. This protein phosphorylates itself in response to binding dsRNA and a key component of the cap-binding complex, eIF2A, that leads to global decapping (50,51). Exposure of HEK293 cells to either Poly(I:C) treatment or SARS-CoV-2 infection resulted in the accumulation of phospho-PKR and phospho-eIF2α, consistent with prior reports showing increased phosphorylation in response to either condition in humans. It also confirms that in our experimental system, SARS-CoV-2 generates enough dsRNA during replication to trigger PKR activation (52). Again, an opposite response is seen in TBLU-ACE2 cells with neither phospho-PKR nor phospho-eIF2α being detectable in response to Poly(I:C) or SARS-CoV-2 infection (Figure 4E). We then sought to impose a human-like PKR response to dsRNA using ectopic expression of human orthologs in the TBLU-ACE2 cells. When the cells were transfected with a plasmid encoding PKR, phosphorylated PKR could be seen in response to Poly(I:C) but not SARS-CoV-2. This is consistent with the responses we observed at the transcript level with dsRNA sensors in the bat cells. Despite the activation of PKR in response to Poly(I:C), there was no activation of bat eIF2α. Upon transfection of eIF2α plasmids, robust phosphorylation of both PKR and eIF2α was seen in response to Poly(I:C) and SARS-CoV-2. Thus, it appears that not only do bats have a block in the sensing of dsRNA by PKR. It is also possible there is an inability of bat PKR to modify the key residue seen in humans of the bat eIF2α ortholog. However, while our results show that the Phospho-PKR antibodies used in these experiments cross react with Bat PKR, this may not be the case for phosphor-eIF2α detection. *T. brasilensis* PKR Thr446 is substituted by Serine (Ser), while Thr451 remains intact (Supp Figure 3). While Ser51 is conserved in *T. brasiliensis*, protein alignment with the human ortholog reveals other adjacent amino acid changes that could interfere with antigen recognition (Supp Figure 4).(53)

Lastly, we evaluated the effects of activation human-like PKR responses by testing cytotoxicity in TBLU-ACE2 and HEK cells after Poly(I:C) treatment (Figure 4F). Control TBLU-ACE2 cells showed no significant increase in cell death when exposed to Poly(I:C). However, TBLU-ACE2 cells transfected with human PKR or eIF2α and treated with Poly(I:C) showed elevated cytotoxicity relative to untreated cells. The increase in cell death was similar after Poly(I:C) treatment in both PKR and eIF2α transfections, suggesting that there may be some other PKR target residues in bats that aren’t the typical site-modified in human eIF2α. HEK293 cells treated with Poly(I:C) displayed severe cytotoxicity, highlighting heightened sensitivity to dsRNA in humans relative to bats (Figure 4E). Indeed, PKR is linked to COVID-19 severity (53–56). These results confirm bat cells bypass the PKR-eIF2α pathway and rely on alternative methods to limit the impacts of viral activity, such as through the involvement of Dicer activity. This highlights a fundamental difference in the bat response to SARS-CoV-2 that could be related to their ability to tolerate viral infections without the harmful effects observed in other species, such as humans.

## Materials and Methods

### Ethics Statement and Biosafety

All the experiments involving live Severe Acute Respiratory Syndrome Coronavirus 2 (SARS-CoV-2, strain USA_WA1/2020; GenBank accession number MT020880) were conducted by certified personnel within USDA-certified Biosafety Level 3 (BSL-3) facilities at the University of Southern Mississippi (USM). These procedures adhered strictly to the biosafety protocol approved by the USM Institutional Biosafety Committee.

### Viruses

The SARS-CoV-2 was obtained from the World Reference Center for Emerging Viruses and Arboviruses at the University of Texas Medical Branch. The parental virus underwent a single passage in Vero E6 cells (catalogue no. CRL-1586, ATCC), after which the viral stock was collected. The viral titer of these stocks was quantified using plaque assays and expressed in plaque-forming units (PFU) per milliliter as described in previous studies (57).

### Cell Culture

The lung epithelial bat cell line TBLU (catalogue no. CCL-88; ATCC), HEK293 (catalogue no. CRL-1573; ATCC), and Vero E6 (catalogue no. CRL-1586, ATCC) was cultured in DMEM:F-12 (Gibco) supplemented with 10% FBS and Penicillin streptomycin. Cryopreserved TBLU and Vero E6 cell lines were successfully resuscitated following standard thawing protocols. All cells were cultured at 37 °C in 5% CO_2_ with the regular passage of every 2–3 d. TBLU-ACE2 stable cell lines overexpressing ACE2 orthologues were maintained in growth medium supplemented with 1 μg ml^−1^ of puromycin. TBLU cells overexpressing the human ACE2 ortholog was generated by transfection of a vector carrying ACE2 coding sequences and the blue fluorescence marker (Addgene plasmid no. 164219) into TBLU cells through Lipofectamine 3000 (Thermo Fisher Scientific). Stable cells expressing various ACE2 orthologues were selected for and maintained in growth medium with puromycin (1 μg ml^−1^). Cells selected for at least 14 d were stable cell lines and used in different experiments. Infections of TBLU and Vero E6 cells used SARS-CoV-2 at MOI of 0.5 and 1.0 when the cells reached 60% confluence. Following a 1-hour incubation period to allow for viral entry, the infection media was replaced with fresh complete media, and the cells were incubated for an additional 48 hours. At 48 hours post-infection, total RNA was extracted and further testing carried out. Cell viability assay was as described in previous studies (58,59).

### Immunofluorescence Assay to evaluate SARS-CoV-2 infection

TBLU, TBLU-hACE2 and Vero E6 cells (100,000 cells per well) were seeded into 24-well glass-bottom plates for overnight incubation and infected with SARS-CoV-2 at 0.5 multiplicity of infection (MOI) for 48 hours. Following infection, cells were fixed with 4% paraformaldehyde, permeabilized with 0.1% Triton X, and blocked with antibody dilution buffer. The cells were stained with a primary SARS-CoV-2 nucleocapsid monoclonal antibody (2 mg mL^-1^, 1 : 500 in ADB, Invitrogen) overnight at 4 °C covered in foil, followed by stained with a FITC conjugated goat anti-rabbit IgG (H + L) cross-adsorbed secondary antibody (2 mg mL^-1^, 1.3 : 1000 in ADB, Invitrogen). The nuclei were stained with 4,6-diamidino-2-phenylindole (DAPI, 300nM, Invitrogen) for 5 minutes at room temperature. Images were then acquired using a Stellaris STED confocal microscope (Leica). This protocol was adapted from a previously published method (57,60–62).

### PCR Quantification of SARS-CoV-2 Nucleocapsid and Dicer expression

TBLU, TBLU-hACE2 and Vero E6 cells were seeded at a density of 5 × 10^5^ cells per well into 6-well plates and incubated overnight. Cells were infected with SARS-CoV-2 at 0.5 MOI for 48 hours. SARS-CoV-2 nucleocapsid and Dicer expression levels of TBLU, TBLU-hACE2 and Vero E6 cells were determined by RT–qPCR. In general, total RNA from cells was extracted by trizol. cDNA was reverse transcribed from 1 μg of total RNA by the Maxima H Minus First Strand cDNA Synthesis Kit (Thermo Scientific) with dsDNase (catalogue no. FERK2561); 1/20 volume of the cDNA was used as the template for the qPCR assay using the Applied Biosystems Fast SYBR Green Master Mix (catalogue no. 4385610) and a CFX96 Touch Real-Time PCR instrument (Bio-Rad Laboratories). Ribosomal RNA was used as an internal control for the normalization of the SARS-CoV-2 relative expression level. Data are presented as the mean ± s.e.m. (*n* = 3). <u>For amplification of SARS-CoV-2 sub-genomic RNAs by RT-PCR,</u> reactions were performed with Phire RT-PCR master mix using primers designed for the target SARS-CoV-2 sub-genomic RNA (spike, nucleocapsid, matrix, envelope) (Supp Figure 5). Controls included uninfected cells with visualization of PCR products via gel electrophoresis.

### Western Blot

Differentially expressed proteins were measured by western blot using extracted proteins from different cell lines and conditions quantified using BCA assay (Thermo Fisher Scientific) method. Proteins were separated by 8–15% SDS-polyacrylamide gel electrophoresis (SDS-PAGE) and transferred to a poly(vinylidene fluoride) (PVDF) membrane (immunobilon-P, 0.45 mm; Millipore, Billerica, MA) using the semidry transfer system (Atto Corp., Tokyo, Japan). The membranes were blocked with 5% low fat milk in Tris-buffered saline containing 1% Tween 20 (TBS-T, pH 7.4) at RT for 1 h and incubated overnight at 4 °C with a 1:1000 dilution of the respective primary antibody (Supp Figure 6). The membranes were washed five times with TBS-T for 5 min each at RT and incubated with a 1:1000–1:5000 dilution of horseradish peroxidase (HRP)-conjugated secondary antibody for 1 hr at RT. The membranes were then washed again five times with TBS-T. The proteins were detected by ECL reagent (Bio-Rad, Hercules, CA) and analyzed using Image Lab 4.1 (Bio-Rad) program. The densitometry readings of the bands were normalized according to Actin expression as control.

### Small RNA sequencing analysis

Total RNA was extracted using Trizol extraction and samples and sequenced by a commercial vendor. Small RNA libraries were constructed using the TruSeq small RNA Library Preparation kit and single-end sequencing was performed to produce 50-bp reads (SE50) using the Illumina NovaSeq platform. After sequencing, adapters are removed using Trim Galore (63). Clean reads are then aligned to the reference genome using Bowtie, with careful handling of mismatches and multimapping reads (64).

The *SARS-CoV-2* (GCF_009858895.2) and *T. brasilensis* (GCA_004025005.1) genomes were acquired from the National Center for Biotechnology Information (NCBI). Small RNA analysis was carried out using pipelines diagrammed in supplement and protocol adapted from a previously published method (65,66). Small RNA loci were identified by aligning reads with Bowtie, converting the alignments to BedGraph format, and applying coverage-based filtering using AWK. Likewise, size distribution was assessed by utilizing AWK to calculate the read lengths from alignments mapped to individual loci.

Fastqs of combined subsets (17–19 nt, 20–22 nt, and 23–24 nt) were aligned to the *SARS-CoV-2* genome (GCF_009858895.2) using Bowtie, with alignments filtered and sorted into BAM files via SAMtools (67). Read coverage was computed using BEDTools to generate BedGraph files (66). Strand-specific BAM files were created to differentiate positive and negative strand reads, and strand-specific coverage data were normalized to reads per million (RPM) and log-transformed using custom scripts.

To analyze vsiRNA overlaps across RNA lengths (15–32 nt), read overlaps were calculated using a publicly available python script (66). For 15-nt RNAs, overlaps with target sequences (15–32 nt) were computed, generating separate FASTA files for each target length. These were parsed to count sequences, and the results consolidated into summary files. This procedure was repeated for all RNA lengths up to 32 nt, adjusting parameters as necessary. The summary data were combined into a single dataset and visualized using a custom R script, yielding a matrix representation of the results.

miRNAs in *T. brasilensis* were analyzed using miRDeep2 with standard parameters (68). Annotations from miRBase were utilized to guide the identification process. Known miRNA candidates called by miRDeep2 were filtered based on a threshold score >50, and loci meeting this criterion were classified as confident. The miRDeep2 analysis provided outputs for both known and novel miRNAs identified in the bat genome (Supplementary file 3, 9).

Heatmap analysis of small RNA (sRNA) bin counts from uninfected datasets were performed using the R package ‘pheatmap’ (69). Rows were scaled, column clustering was disabled, and hierarchical clustering was applied to identify distinct groups. Row order from clustering was extracted and mapped to the infected libraries for direct comparison, revealing distinct small RNA expression profiles between conditions.

### Computational Pipeline for mRNAseq

Sequencing was carried out a commercial vendor, and raw reads were quality-checked using FastQC. The *T. brasilensis* genome was indexed with STAR, and paired-end RNA-Seq data were aligned to the reference genome. Gene-level read counts were quantified, and the aligned data were saved in a sorted BAM file along with related output files. Genomic annotation was used to quantify mRNA transcripts from SARS-CoV-2-infected and uninfected TBLU-ACE2 cells using the “GCA_030848825.1_DD_mTadBra1_pri.xenoRefGene.gtf” file from the UCSC Genome Browser. RNA-Seq data were analyzed with the DESeq2 package in R to assess differential gene expression across experimental conditions (70). The count and metadata files were imported and stored in an object alongside the design formula that incorporated both Treatment and Infection factors. Genes with low expression were filtered out by retaining only those with a sum of counts greater than 1 across all samples. Variance stabilizing transformation (VST) was applied for normalization, with the ‘blind = FALS’ parameter set to retain the structure of the data according to the experimental design, preserving biological variation between conditions (treatment vs. control) (Supp Figure 2). Differential expression analysis was conducted for specific contrasts, and the results were visualized using MA plots and saved to text files. For Gene Ontology (GO) analysis the TopGO R package was used (71). GO terms associated with the RefSeq mRNA accessions of GCA_030848825 xenoRefGene were retrieved using the **bio**logical **D**ata**B**ase **net**work (**bioDBnet**) tool (72).

## RESULTS

### DISCUSSION

SARS-CoV-2 RNAs appear to be processed by Dicer in bats based on the presence of vsiRNA fragments that are lost upon depletion of Dicer. Decreasing Dicer activity also leads to higher rates of viral replication, suggesting that the processing of genomic RNAs by Dicer is a component of the bat cell antiviral response. Infection of Vero E6 cells with SARS-CoV-2 shows a much greater viral load than the TBLU cell line used in these studies, showing a different and more effective response in bat cells. A significant aspect of the anti-viral response in bats appears to be related to their handling of dsRNA accumulation in cells where treatment with Poly(I:C) leads to low cytotoxicity compared to human cells. Indeed, introduction of the human dsRNA sensor, PKR, and its target substrate significantly increases cell death. Remodeling the transcriptome in response to SARS-CoV-2 infection includes alteration in endogenous small RNA processing. The interplay of these adaptations, such as dampened PKR response, avoids translational arrest and inflammation response could contribute to the ability of bats to endure high viral loads and coexist with lethal RNA viruses like SARS-CoV-2 without developing disease.

In many eukaryotes, Dicer performs two functions: regulating endogenous genes through the generation of miRNAs and mediating the antiviral response through vsiRNA production (38). Other species of small RNA are recognized Dicer products; however, they are often species-specific with cryptic functions (73). Many animals possess multiple Dicer proteins, with a dedicated miRNA-Dicer homolog and a separate gene encoding a siRNA-Dicer that has a role in antiviral immunity (74). Unlike these organisms, vertebrates have a single Dicer gene that processes miRNAs. The loss of anti-viral Dicer appears to have occurred very early in the deuterostomes, being replaced by alternative antiviral mechanisms like interferon and cell-based immunity (38). Our results show that convergent evolution has occurred with bat Dicer, which has a reactivated antiviral function. This is consistent with a study using bat cells from a different species, *Pteropus Alecto* (47)*. P. Alecto* is an old-world fruit bat, while the origin of the cells used in this work is from *T. brasilensis,* a new-world bat (75). The presumptive vector of SARS-Cov-2 is a Phinolophid, which are more closely related to *P Alecto.* Our work shows that the reactivated antiviral response of Dicer occurred very early in the evolution of bat lineages possibly coinciding with the time flight evolved in these mammals. This suggests a linkage to the evolution of flight physiology and a requirement for anti-viral mechanisms disconnected from the febrile response.

This work also suggests that bat-originating viruses, such as SARS-CoV-2, will likely have evolved counter-mechanisms to limit the efficacy of Dicer as an antiviral defense (76,77). Emerging evidence suggests that SARS-CoV-2 nucleocapsid proteins, such as NSP15 and NSP8, play a role in dampening the host’s small RNA-mediated defenses (76). Thus, it is plausible that SARS-CoV-2 may possess mechanisms that interfere with Dicer activity similar to how plant VSRs, such as HcPro, P21, P19, P15, and P0, which disrupt miRNA-in other vertebrate hosts, further impacting pathophysiology of the virus (39–42). Supporting this, it was reported that a significant reduction in Dicer expression in COVID-19 patients was seen compared to healthy individuals (49) Further investigation of SARS-CoV-2 proteins could offer insights into whether the virus manipulates the processing of small RNAs to disrupt Dicer activity.

In humans, SARS-CoV-2 can drive severe pathologies resulting from hyperactivation of an innate immune response which becomes the primary factor contributing to severe complications, including acute respiratory distress syndrome (ARDS), multi-organ damage, and vascular issues (78,79). dsRNA is a major activator of the antiviral response, and its accumulation may be a component of the overreaction of the human immune system to SARS-CoV-2 (80). This raises an interesting question regarding the replication behaviors of SARS-CoV-2 relative to the “domesticated” common-cold coronaviruses HCoV-229E and HCoV-NL63. Superficially common cold coronaviruses share many of the genetic characteristics of the SARS-CoV-2 virus (81,82). All are large positive-strand viruses with similar early and late proteins. During infection, both deform endo-membranes to form replication sites (83,84). How do the common cold viruses avoid activating the dsRNA response, which may be a component of the reaction to SARS-CoV-2? Understanding the alterations in viral genes that impact the presentation of dsRNA to cytoplasmic proteins could be an aspect of the vastly more benign behavior of common cold coronaviruses. Targeting dsRNA-generating processes in SARS-CoV-2 or other deadly coronaviruses could yield a novel approach to limiting immune-related damage associated with these infections.

### CONCLUSION

SARS-CoV-2 RNAs appear to be processed by Dicer in bats, as evidenced by the presence of vsiRNA fragments, which are diminished upon Dicer depletion. Reduced Dicer activity correlates with increased viral replication, suggesting that Dicer-mediated processing of genomic RNAs is a key component of the bat antiviral response. Consequently, bats can handle dsRNA accumulation with minimal cytotoxicity, as demonstrated by the low toxicity of Poly(I:C) treatment in bat cells compared to human cells. SARS-CoV-2 infection in bats also prompts transcriptome remodeling, including changes in endogenous small RNA processing. Adaptations such as a dampened PKR response help avoid translational arrest and limit inflammatory responses, enabling bats to tolerate high viral loads and coexist with lethal RNA viruses like SARS-CoV-2 without succumbing to disease. An important implication of this study is the potential for targeting dsRNA-generating processes in SARS-CoV-2 and other deadly coronaviruses to mitigate immune-related damage. This strategy could offer a novel approach to reducing the pathological consequences of these infections in humans.

## LIST OF ABBREVIATIONS

siRNA: Small-interfering RNA
OAS1: 2’-5’-Oligoadenylate Synthetase 1
MX1: Myxovirus Resistance Protein 1
dsRNA: Double-stranded RNA
PKR: Protein Kinase R
SARS-CoV-2: Severe Acute Respiratory Syndrome Coronavirus 2
miRNA: microRNA
SARS-CoV: Severe Acute Respiratory Syndrome Coronavirus
MERS-CoV: Middle East Respiratory Syndrome Coronavirus
ACE2: Angiotensin-Converting Enzyme 2
Spike (S): Spike Protein
Envelope (E): Envelope Protein
Matrix (M): Matrix Protein
Nucleocapsid (N): Nucleocapsid Protein
nsp: Non-structural Protein
HCoV-229E: Human Coronavirus 229E
HCoV-NL63: Human Coronavirus NL63
P. alecto: Pteropus alecto
IFN: Type I and Type II Interferon
NF-κB: Nuclear Factor Kappa-light-chain-enhancer of Activated B Cells
NLRP3: NOD-, LRR- and Pyrin Domain-containing Protein 3
TLRs: Toll-like Receptors
eIF2α: Eukaryotic Initiation Factor 2 Alpha
ATF4: Activating Transcription Factor 4
ATF3: Activating Transcription Factor 3
RNAi: RNA Interference
VSRs: Viral Suppressors of RNA Silencing
TBLU: Tadarida brasiliensis (Brazilian Free-tailed Bat)
sgRNAs: Subgenomic RNAs
qPCR: Quantitative Polymerase Chain Reaction
PCR: Polymerase Chain Reaction
DKD: Double Knockdown
MOI: Multiplicity of Infection
Vero E6: Epithelial Cells Derived from African Green Monkey Kidney
293T: Human Embryonic Kidney 293 Cells Expressing SV40 T-antigen
BHK: Baby Hamster Kidney Cells
vsiRNAs: Virus-derived Small-interfering RNAs
SINV: Sindbis Virus
PaKi cells: Pteropus alecto Kidney Cells
mRNA: messenger RNA
GO: Gene Ontology
Poly(I:C): Polyinosinic:polycytidylic acid
FPKM: Fragments Per Kilobase of Transcript Per Million Mapped Reads
HEK293: Human Embryonic Kidney 293 Cells
Phospho: Phosphorylated
Thr: Threonine
Ser: Serine
OB: Oligonucleotide Binding
CTD: C-terminal Domain
P-loop: Phosphate-binding Loop
COVID-19: Coronavirus Disease 2019
BSL-3: Biosafety Level 3
PFU: Plaque Forming Units
RT–qPCR: Reverse Transcription Quantitative Polymerase Chain Reaction
TBS-T: Tris-Buffered Saline with Tween
HRP: Horseradish Peroxidase
PVDF: Polyvinylidene Fluoride
SDS-PAGE: Sodium Dodecyl Sulfate Polyacrylamide Gel Electrophoresis
VST: Variance Stabilizing Transformation

## DECLARATIONS

### 1. Ethics approval and consent to participate

Not Applicable

### 2. Consent for publication

Not Applicable

### 3. Availability of data and materials

The data resulting from High throughput sequencing (fastq files) has been deposited in the Sequence Read Archive (SRA) repository under bioproject number PRJNA1207558.

### 6. Authors’ contributions

**I.O**: contributed to the conceptualization of the study, conducted and implemented experiments, data analysis, manuscript writing, and review.

**S.U**: implemented infection of cell lines with virus and participated in manuscript review.

**S.K**: data gathering and contributed to manuscript review.

**S.V**: conducted experimental work and manuscript review.

**C.F**: conducted experimental work and manuscript review.

**F.B**: SARS-CoV-2 virus, veroE6 cell line, virus infection and manuscript review.

**A.F**: contributed to the conceptualization of the study, implemented experiments, lab and resource, data analysis, manuscript writing, and review.

## 7. Acknowledgements

