## supplementary materials for "Processing of Genomic RNAs by Dicer in Bat Cells Limits SARS-CoV-2 Replication"

**Supplementary Figures**

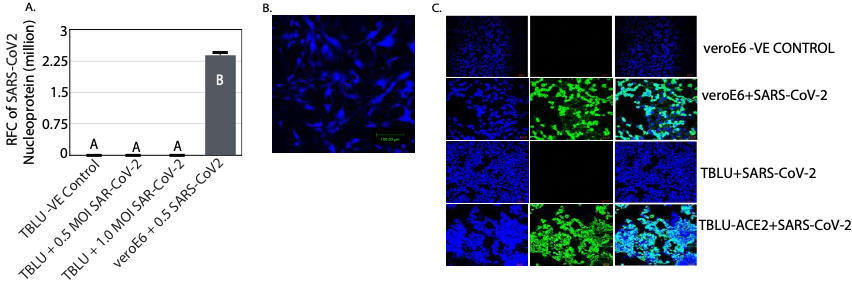

**Figure S1. TBLU challenged with SARS-CoV-2 virus.**

(A) Bar graph depicting SARS-CoV-2 nucleocapsid protein expression in normal TBLU cells. (B) Human ACE2-overexpressing TBLU (TBLU-ACE2) cells visualized with a blue fluorescence reporter. (C) Immunostaining analysis of SARS-CoV-2 nucleocapsid protein expression in VeroE6, TBLU and TBLU-ACE2 cells 48 hours post-infection. Statistical significance was determined using Tukey’s ANOVA.

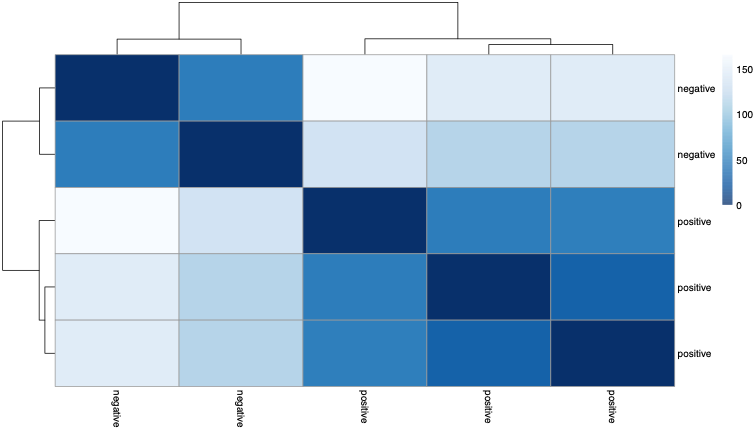

**Figure S2. Quality assessment of mRNA sequencing data.**

(A) VSD plot showing consistent alignment of features across SARS-CoV-2-infected triplicates and uninfected duplicates.

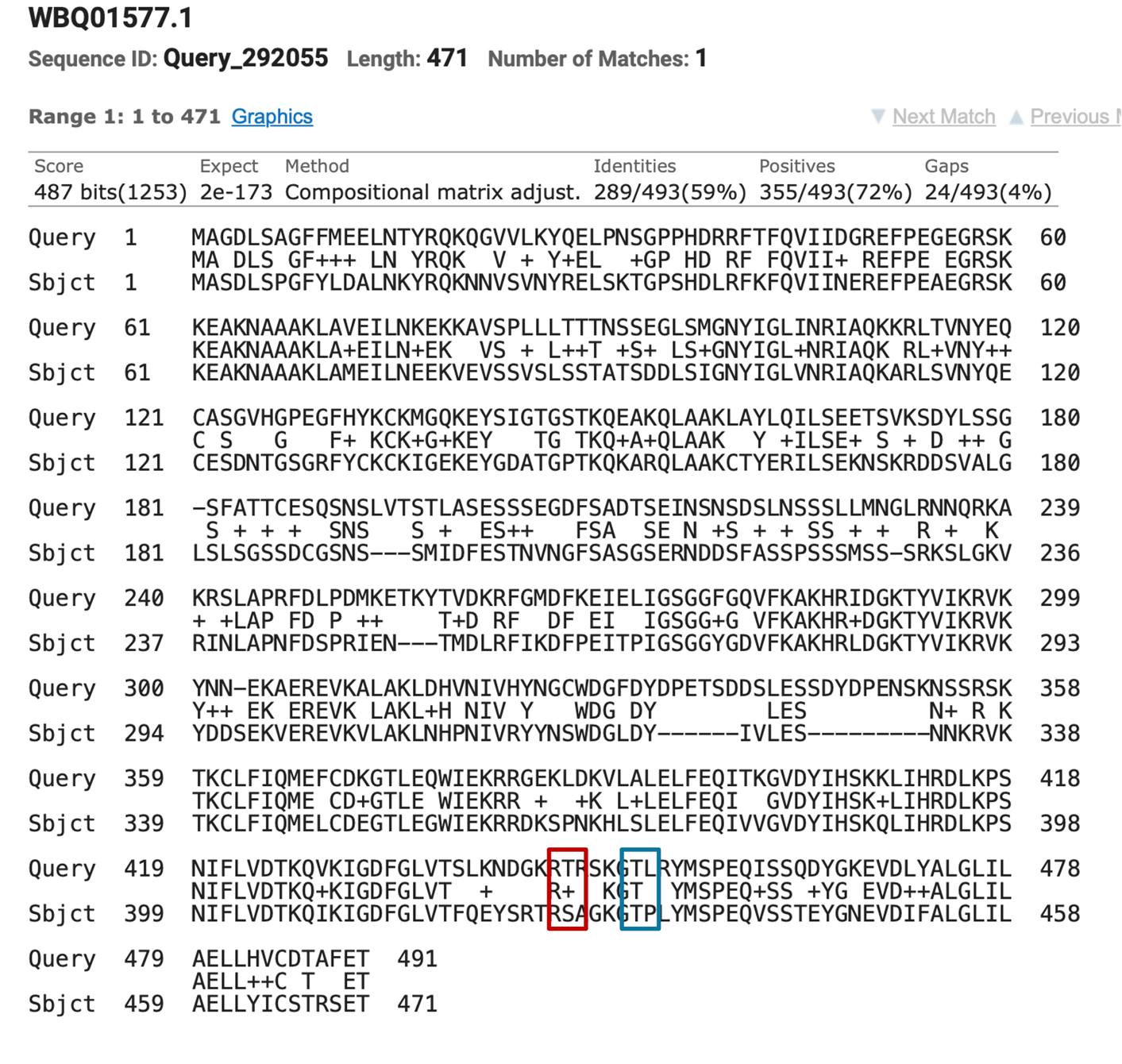
**Figure S3. PKR.** Thr446 and Thr451 are key residues in the activation loop of PKR, and their phosphorylation is crucial for its activation. In bat PKR, Thr446 is substituted by Ser (red box), while Thr451 (blue box) remains unchanged.

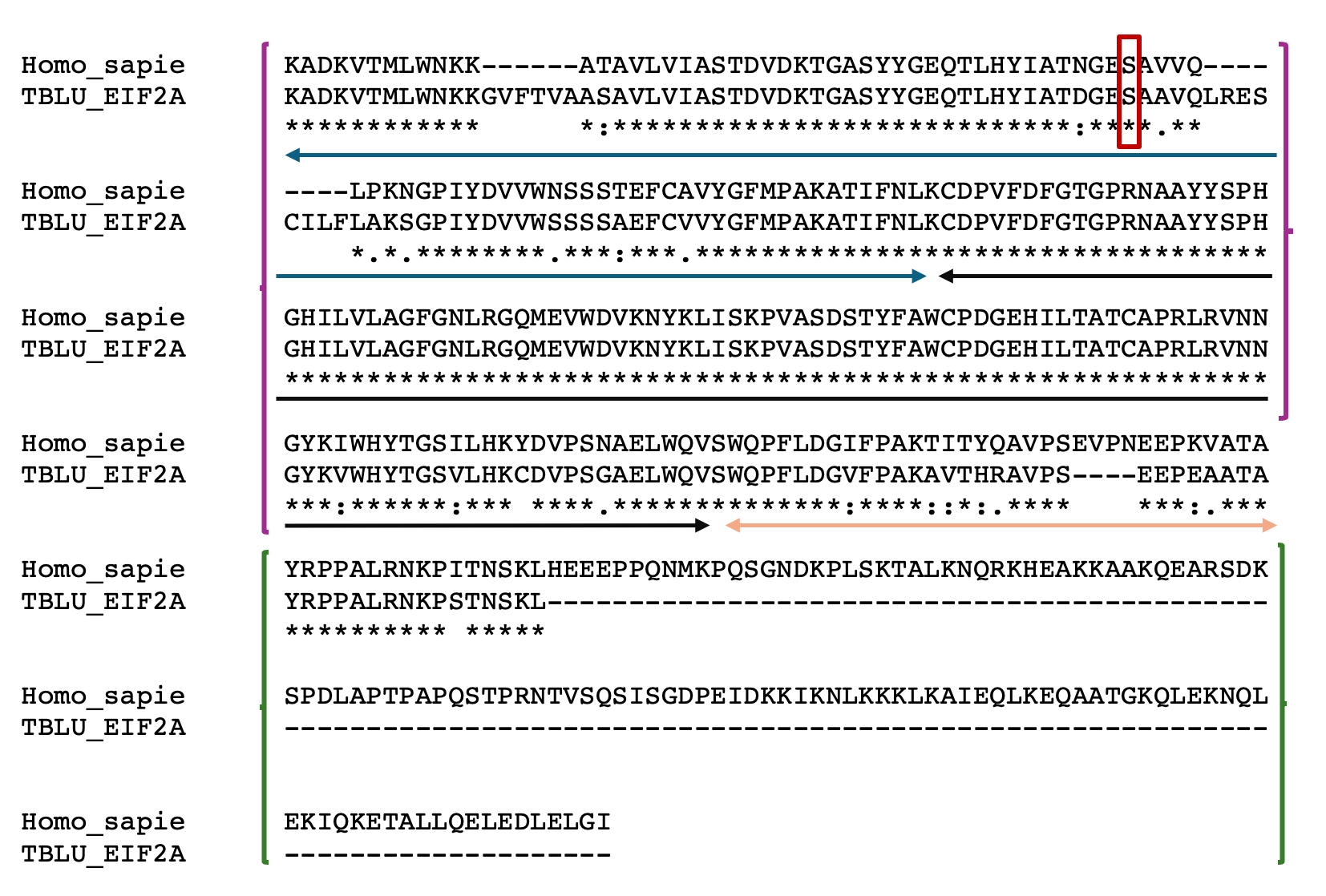

**Figure S4. EIF2A.**

Using the MUSCLE tool on phylogeny.fr, we aligned the protein sequences of human and bat eIF2α to compare structural features. The structure of human eIF2α comprises two folded domains: an N-terminal domain (NTD, residues 1–183) (Purple open-close box) and a C-terminal domain (CTD, residues 188–280) (Green open-close box), connected by a flexible linker (Orange arrow). The NTD includes two subdomains: an oligonucleotide/oligosaccharide-binding (OB)-fold subdomain (residues 18–85) (blue arrow), containing the phosphorylation loop (P-loop) with the regulatory Ser51 site (red box), and an α-helical subdomain (residues 91–183) (black arrow). Our analysis revealed several key differences in bat eIF2α: the OB-fold subdomain contains insertions that could alter the structure of the P-loop, the α-helical subdomain shows minor substitutions, and most of the CTD appears to be intrinsically disordered. These structural differences may have functional implications for eIF2α-mediated translation regulation in bats.

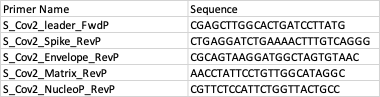

**Figure S5. Primers.** Primers designed for SARS-CoV-2 sub-genomic RNA amplicification.

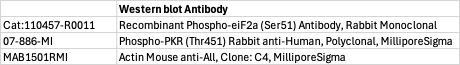

**Figure S6. Antibody.** Antibodies used for western blot.
